# Impact Evaluation of Significant Feature Set in Cross Project for Defect Prediction through Hybrid Feature Selection in Multiclass

**DOI:** 10.1101/2023.07.20.549868

**Authors:** Sana Gul, Rizwan Bin Faiz, Mohammad Aljaidi, Ghassan Samara, Ayoub Alsarhan, Ahmad al-Qerem

## Abstract

Cross-project defect prediction (CPDP) is a significant way of defect identification in the project. In cross-project defect prediction, we extract knowledge from the source project and apply that learned knowledge to predict labels for the target project. However, the model performance can be affected by features that are insignificant and irrelevant. Hybrid feature selection (HFS) can play a significant role in achieving high prediction accuracy by selecting significant and only relevant features. Our aim is to explore effect of significant feature selection through a hybrid approach upon cross-project (CP) defect prediction for datasets which are multi-class in nature. We leveraged the strengths of Random Forest (RF) and Recursive Feature Elimination Cross Validation (RFECV) which can constructively select few features which are significant. The design of our controlled experiment is 1 Factor 2 Treatments (1F2T). Exploratory Data Analysis (EDA) proves that all versions of PROMISE repository are multi class and have duplicated rows in data, distribution gap among values, and imbalance classes. Hence after removing duplicated rows, reducing the gap present in distribution of data, and balancing classes, we selected significant feature set through Hybrid approach i.e. Random Forest (RF) and Recursive Feature Elimination Cross Validation (RFECV). We used Convolutional Neural Network (CNN) as a classifier to predict Cross project defects along with SoftMax as the last layer. Our experimental setup resulted in the average 78% prediction accuracy measure of all 14 versions in terms of AUC. Our experimental result showed that there is significant impact of HFS on defect prediction accuracy for different datasets in the CP.

## 1. Introduction

In today’s era, software is becoming more and more complex, increasing the significance of software reliability. Despite rigorous testing of software, some defects remain in software which leads to unexpected system failure after deployment resulting in a financial loss that the organization has to bear [1][2]. Bad design and code implementation are significant reasons for these defects in software [3]. To make reliable software it is important to do a significant amount of debugging and testing to reduce cost and time and increase productivity. Software defect prediction (SDP) techniques can search defects in software accurately at the beginning of software engineering which reduces cost and time being spent on testing [4]. A defect prediction model can efficiently find the defects in modules by spending less amount of software development and maintenance resources [5].

Many researchers worked on CPDP but achieving a satisfactory result is quite challenging and improvement can be made in this area of research. In CDPD, we come across significant issues of noise, gap in distribution of data between source and target datasets [6], which breach the basic requirement of modeling technology, and also class imbalance issues among two different datasets that negatively impact of the accuracy of CDPD. PROMISE repository being used in this research is a commonly used dataset for software defect prediction studies [7]. This repository is multi-class and also contains noise, data distribution gaps, and class imbalance issues. To solve the issue of the gap in distribution of data, min-max normalization is used and CTGAN synthesizer for balancing the imbalance classes by generating synthetic data for tabular data.

The dataset used for defect prediction consists of features that are used to train the model to predict defects in target software. However, all features are not helpful in the prediction accuracy of the model. Dataset may have a presence of some irrelevant and insignificant features which critically decrease the performance of the prediction model and increase the computational cost of the model as well [8]. Therefore, it is crucial to select the significant and relevant feature set which effectively contribute to software defect prediction. The HFS technique can be beneficial as it can leverage the strength of algorithms and produce better accuracy for defect prediction.

Previous research [8] showed that the model’s performance depends on the factor of relevant features and the classifier being used. It motivated us to find a powerful classifier with superior performance. In recent years, many studies [9] [10] [11] have proposed to extract hidden semantics through deep learning. It is evident form literature that the pooling layer of the convolutional neural network has the capability of outperforming classification results along with SoftMax as a final layer. This layer gives probability of each output of class from 0 to 1. For each particular instance class with highest probability is selected as an output. Also, SoftMax is used for multi-classification which is more suitable for multi-class datasets [12]. Due to its simplicity and probabilistic interpretation, the SoftMax function is generally adopted by many CNN models [13].

The **main contributions** of this paper are as follows:

- We considered PROMISE repository (2021) as multi-class as evident from EDA and then preprocessed e.g. removed noise, minimized data distribution gap, and then balanced classes.
- We then explore the impact of significant feature set upon classification for multiclass in CPDP. We used Wrapper and Embedded methods as hybrid approach i.e. Random Forest combined with Recursive feature elimination Cross Validation algorithms which is highly effective in high-dimensional feature spaces for significant selection from PROMISE repository (2021) for multi-class problem. Then defects are predicted in CP through deep neural network i.e. CNN classifier along with SoftMax as last layer.

## 2. Literature Review

CPDP has become point of concern in literature. Many researchers constructed unique CPDP models to predict defects in datasets while considering all features of the dataset However, achieving high prediction accuracy in CDPD has been a challenge especially when all features regardless of their insignificance. Many studies reviewed in the literature have used non-hybrid feature selection algorithms to select significant features and resulted in improved accuracy of software defect prediction models.

[15] Predicted defects in cross project from PROMISE repository as multi-class. Before using the dataset, noise and imbalance issues in classes of datasets were resolved in the dataset. They have used all features from the dataset for software defect prediction regardless of their insignificance. Their prediction accuracy was 0.7034 in F-measure and 0.6944 in G-measure.

[14] Predicted defects in cross project from PROMISE repository as binary class. They used the InfoGain algorithm as a non-hybrid technique for selection of features. They used Navies Bayes for classification in experiment. They used Precision, Recall, and AUC to measure the result of experiment.

[16] Predicted defects in cross project from PROMISE repository as binary class. They removed duplicated row in data, gap in distribution of data, and imbalance of class’s issues in the dataset. They performed experiments on all features of the dataset and used the CNN model for the prediction of defects in the dataset. Effectiveness of their approach was demonstrated F-measure being 0.66.

[17] Predicted defects in cross project from PROMISE repository as binary class. They have used all features of the dataset for software defect prediction and the CNN model for classification. F-measure was used to measure the results of experiment.

[8] Researchers worked on defect prediction using PROMISE, RELINK, and AEEEM repositories.

They resolved data distribution and class Imbalance issues in the dataset. They used Whale Optimization and Simulated Annealing algorithms as a HFS to select significant features from the dataset. They used the CNN model with KELM as a hybrid neural network for defect prediction. Although they achieved high prediction accuracy through HFS, they did not consider the dataset as multi-class which is evident through EDA.

[18] Predicted defects in cross project from PROMISE repository as multi-class. They used RF and RFECV as HFS approach to select significant features from the dataset. They used the NN layer model as a classifier for the prediction. In the end, experimental results were measured in AUC.

[19] Predicted defects in cross project from PROMISE repository as multi-class. They resolved noise, data distribution gaps, and class imbalance issues in the dataset. They used XGBoost as a non-hybrid approach for the selection of significant features from the dataset. They used CNN model for classification. The effectiveness of their approach was demonstrated through AUC being 75.57%.

[20] Worked on Covid detection of patient using the COVID-19 dataset. They used Filter methods and Genetic Algorithm as HFS approach to select most significant features from the Chest Computed Tomography (CT) images for COVID-19 patients and non-covid patients as a non-hybrid approach for the selection of significant features from the dataset. They used enhanced KNN model for classification. Effectiveness of their approach was demonstrated through F-measure being 93%

[21] Worked on human activity recognition using the UCI machine learning repository and treated it as a multi-class problem. They used Sequential floating forward search to select optimal feature and then sub-optimal feature sets are passed into wrapper procedure to where the result of classification accuracies is calculated using SVM for each sub-optimal feature vector. They constructed the experiment, and the result analysis has showed effectiveness of the proposed methodology with 96.81%

[22] Worked on analysis of expression level of genes for cancer classification using the Colon, SRBC, LEUKEMA1, LEUKEMA2, PROSTATE and lymphoma datasets. They compared the results of different HFS approach which are based on Genetic Algorithm (GA), Ant colony Optimization (ACO), Bat Algorithm (BA), Artificial Bee Colony (ABC), Particle Swarm Optimization (PSO), Black Hole Algorithm (BH), Biogeography Algorithm (BA), Harmony Search Algorithm (HSA) and Grass Hooper Optimization Algorithm (GOA). They performed experiment and concluded that Genetic Algorithm (GA) resulted in highest accuracy with relatively small number of selected genes.

[23] Worked on automatic recognition sign language on seven publicly available benchmark datasets which are ISL, MNIST, Jochen Triesch Static Hand Posture, NUS Dataset II, ASL Dataset, IEEE ASL Dataset, and Arabic. They proposed mRMR-PSO methodology in which Minimum Redundancy and Maximum Relevancy and Particle Swarm Optimization are used as hybrid approach for selection of significant features. They used Support Vector Machine (SVM) for classification in multi-class. After experiment they presented accuracies for ISL and NUS dataset is 99.18% and 96.78% respectively.

[24] Worked on malignant disease prediction using five biological datasets. They removed noise and redundant data in the dataset. They used. They used Conditional Mutual Information Maximization and Binary Genetic Algorithm as HFS approach along with numerous classifiers used for classification which are Naïve Bayes, SVM and K-NN. They demonstrated that their proposed method provides additional support to the significant reduction of the features and gives higher prediction accuracy for malignant diseases.

[25] Proposed to solve feature selection problems through hybrid approach using Grey Wolf Optimization and Particle Swarm Optimization on datasets from UCI Irvine repository. They used KNN model with Euclidean separation matric for classification. The effectiveness of their approach was demonstrated through average accuracy being 0.93.

### 2.2 Research Gap

Through critical analysis of the literature review conducted above, we constructed a research gap depending on the elements that most researchers missed while conducting the research. The research gap on basis of the above literature review is explained as follows:

1. It is apparent after EDA that PROMISE dataset (2021) has multi-class nature there are traces of duplicated rows, gap in distribution, and imbalance of class’s issues. We conducted literature review of these papers [8] [14] [15] [16] [17] which pointed out the fact that researchers have resolved duplicated rows, gap in distribution, and imbalance of classes issues in a disassociating manner. However, we are going to resolve these issues in the promise repository as cross-project multi-class data set in an associated manner.
2. [19] Have worked on CPDP for multi-class upon PROMISE repository which was publicly available in 2020.Whereas, we have worked upon CPDP for multi-class using updated PROMISE repository which were publicly available in 2021. The updated repository has following differences from repository which was available in 2020.
3. [8] Has manifested the impact of HFS upon a binary classification problem in CPDP upon PROMISE, ReLink, and AEEEM repositories. However, its impact on multi-class in CPDP is yet to be explored. As we must select significant features from all features in the dataset, Random Forest is an algorithm that easily handles high-dimensional feature spaces for multiclass problems [26].
4. Our research explores impact of deep neural network i.e., CNN classifier along with SoftMax layer after HFS through Recursive feature elimination Cross Validation and Random Forest algorithms for Cross project in multi-class defect prediction which has not yet been explored in literature.
5. [8] [16] [17] literature made evident that the CNN classifier is used on the PROMISE repository as a binary class which helped to achieve higher defect prediction accuracy. Since [17] [27] literature made it evident that SoftMax improves prediction accuracy more than KELM in multi-classification due to intra-class compactness and inter-class separability of features learned by CNN. Therefore, there is a need to evaluate the effect of CNN classification model SoftMax as a last layer for Multi-class in CPDP.

## 3 Materials and Methods

Firstly, we have mentioned the necessary information about the dataset. After that, we presented our proposed approach step by step in Figure 1.

**Figure 1.**
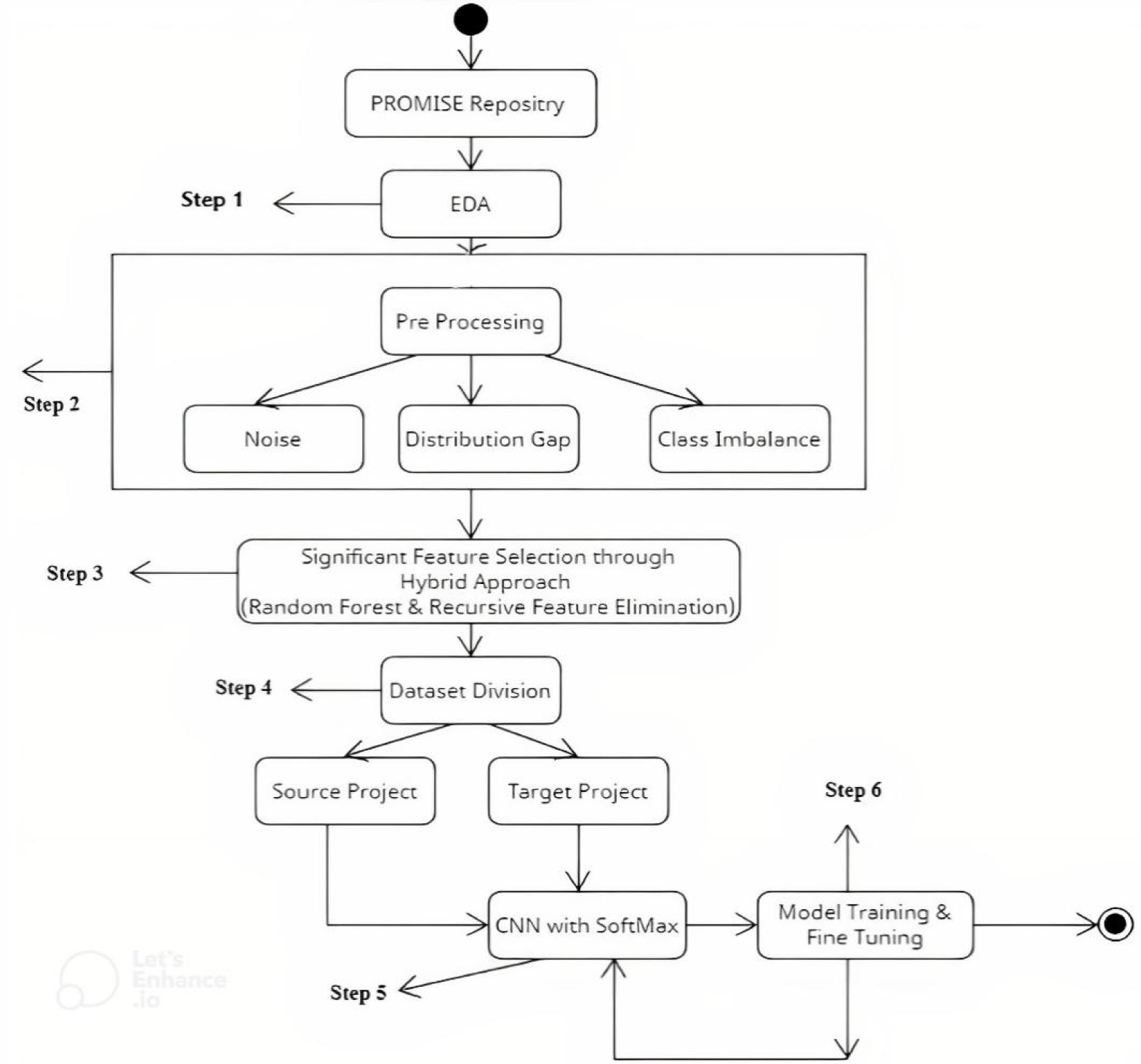
Our proposed methodology

### 3.1 EDA

EDA is performed on PROMISE repository [28] which one of the most widely utilized repository to perform experiment for SDP. The repository’s latest version is published in 2021 and no research has been performed on updated version till now. Some dataset versions which were part of PROMISE repository 2020 are now unavailable in repository as published in 2021 e.g., Forrest 0.7, Forrest 0.8, Ckjm and Xalan-2.4 etc. Also the number of instances in each feature set are different in both repositories. Each dataset consists of 20 features, each having 14 versions. After EDA, it became evident that PROMISE repository has multi-class characteristics instead of binary class having noise, distribution gap, and class imbalance issues. Visual description of the multiple classes of the dataset can better be described in Figures 2 and 3.

**Figure 2.**
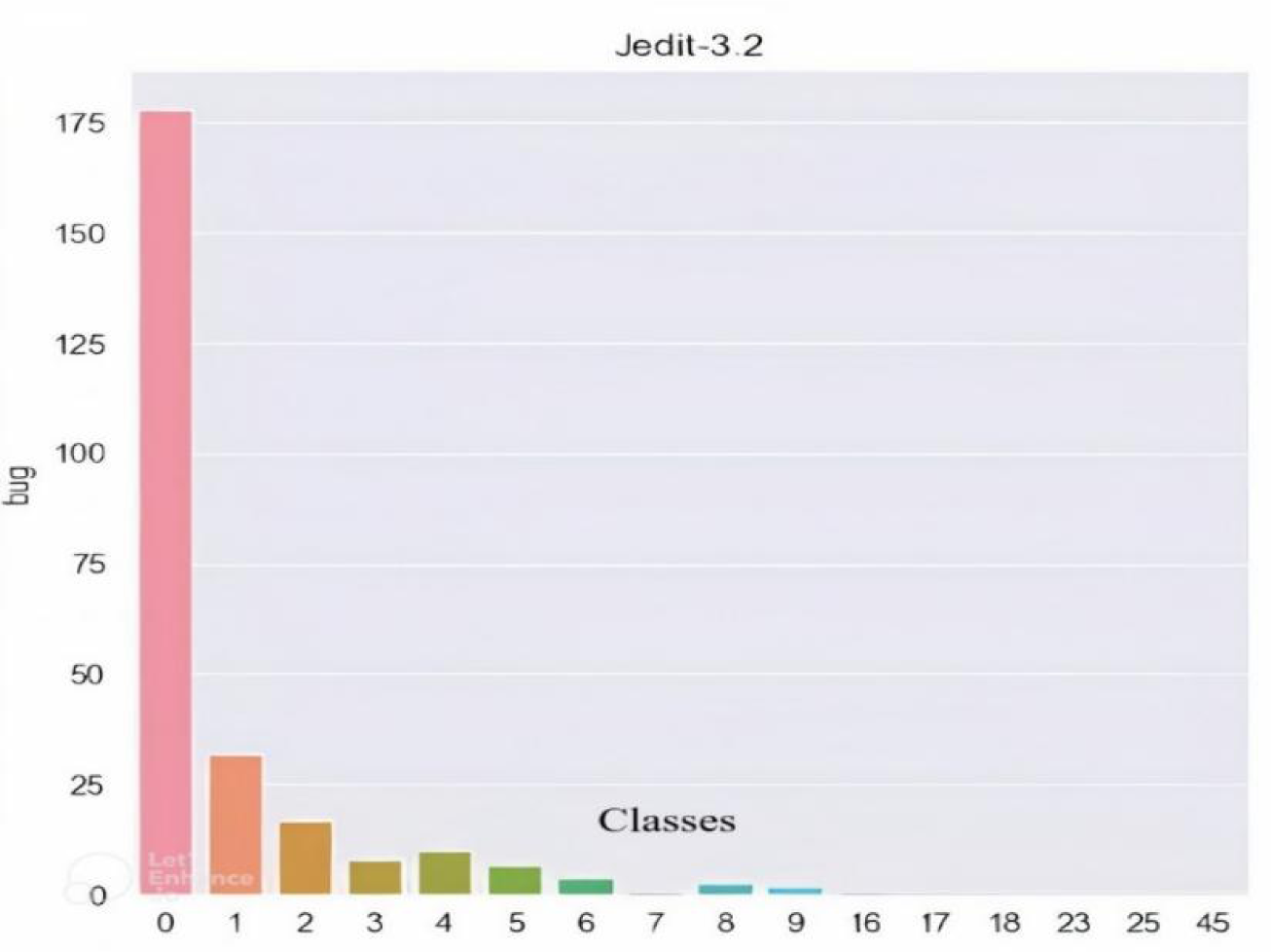
Dataset jedit-3.2 having multiple classes.

**Figure 3.**
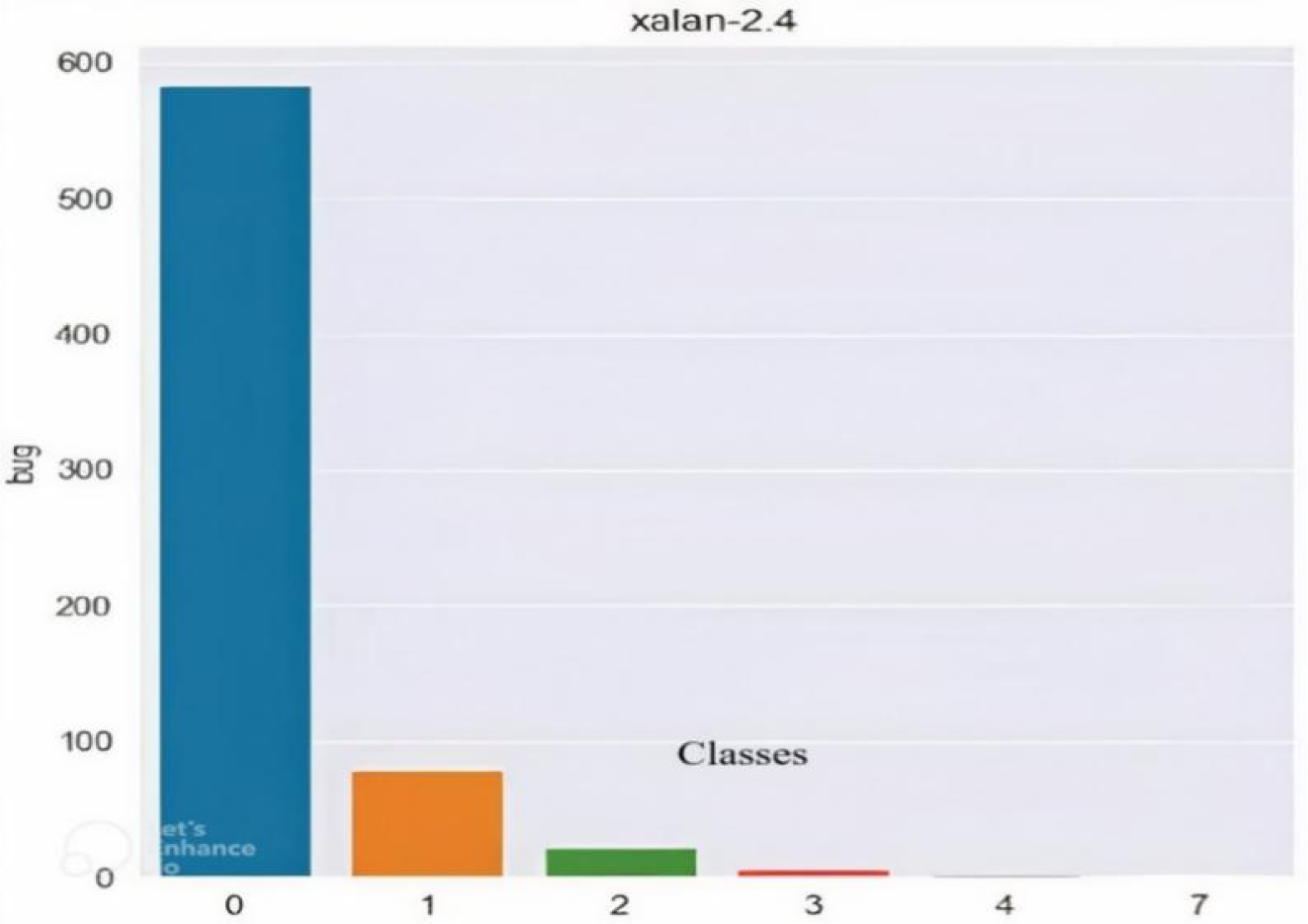
Dataset xalan-2.4 having multiple classes.

**Figure 4.**
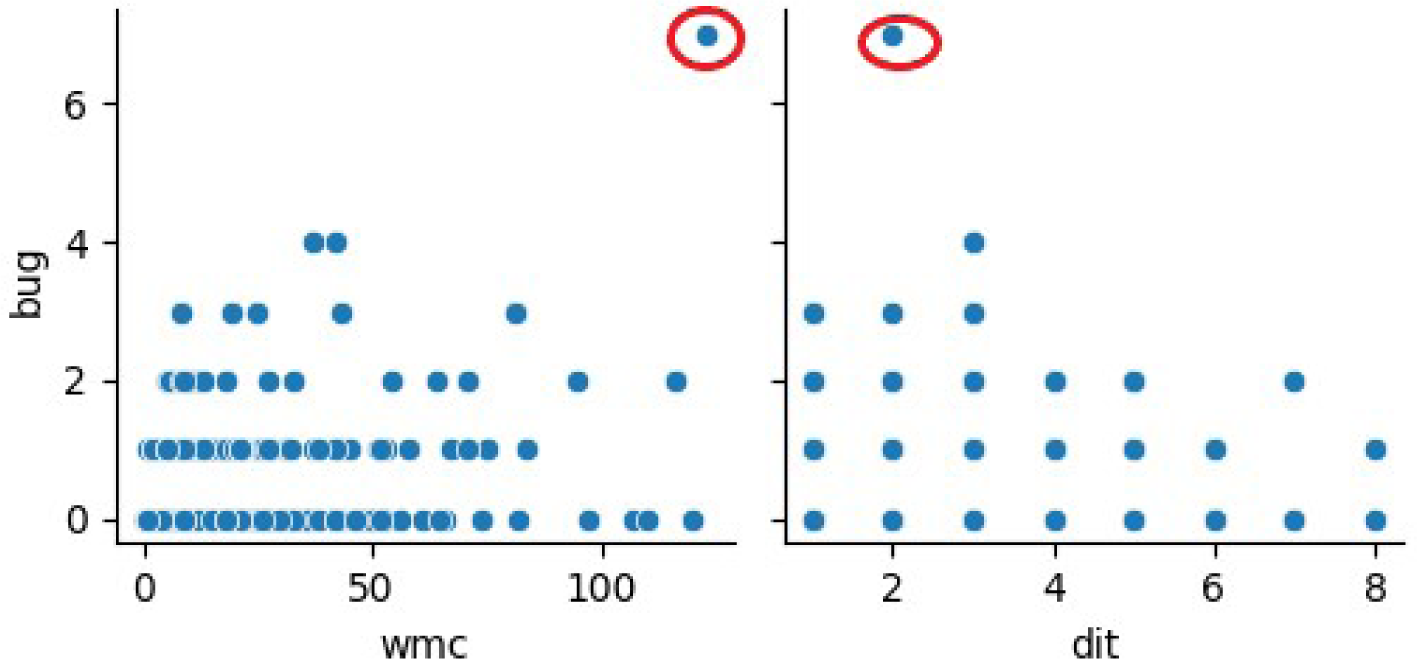
Before removing the gap in distribution of data of the “wmc” Xalan-2.4

**Figure 5.**
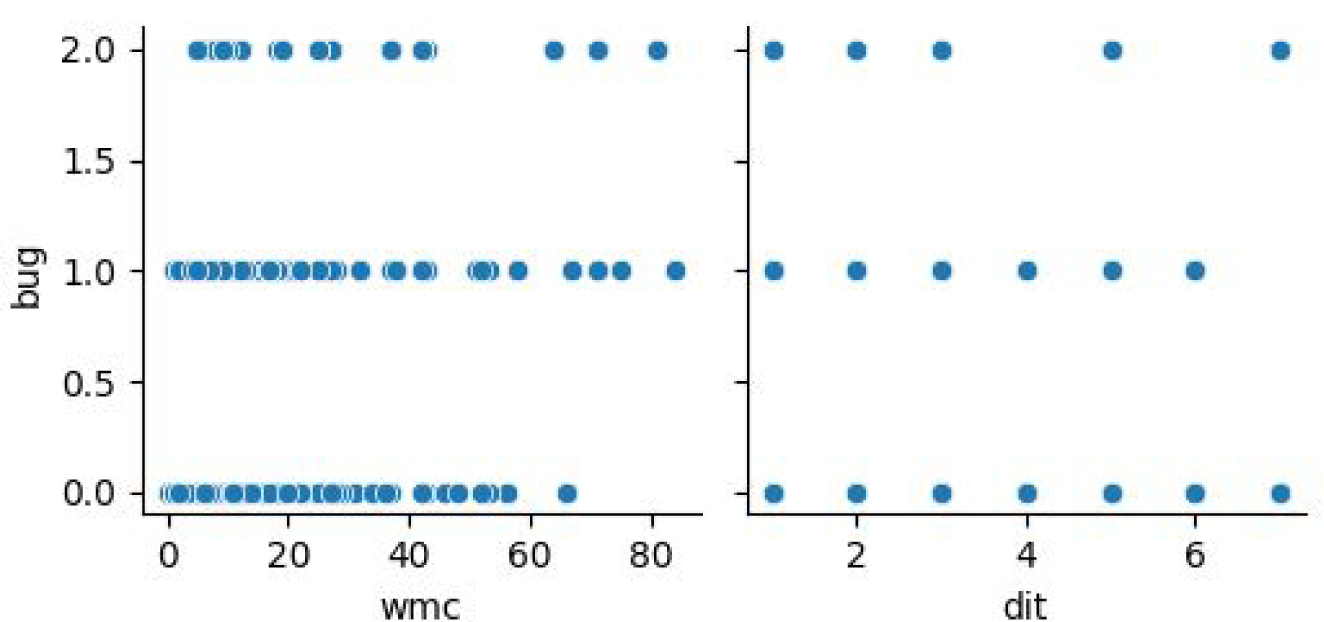
After removing the gap in distribution of data of the “wmc” of xalan-2.4

**Figure 6.**
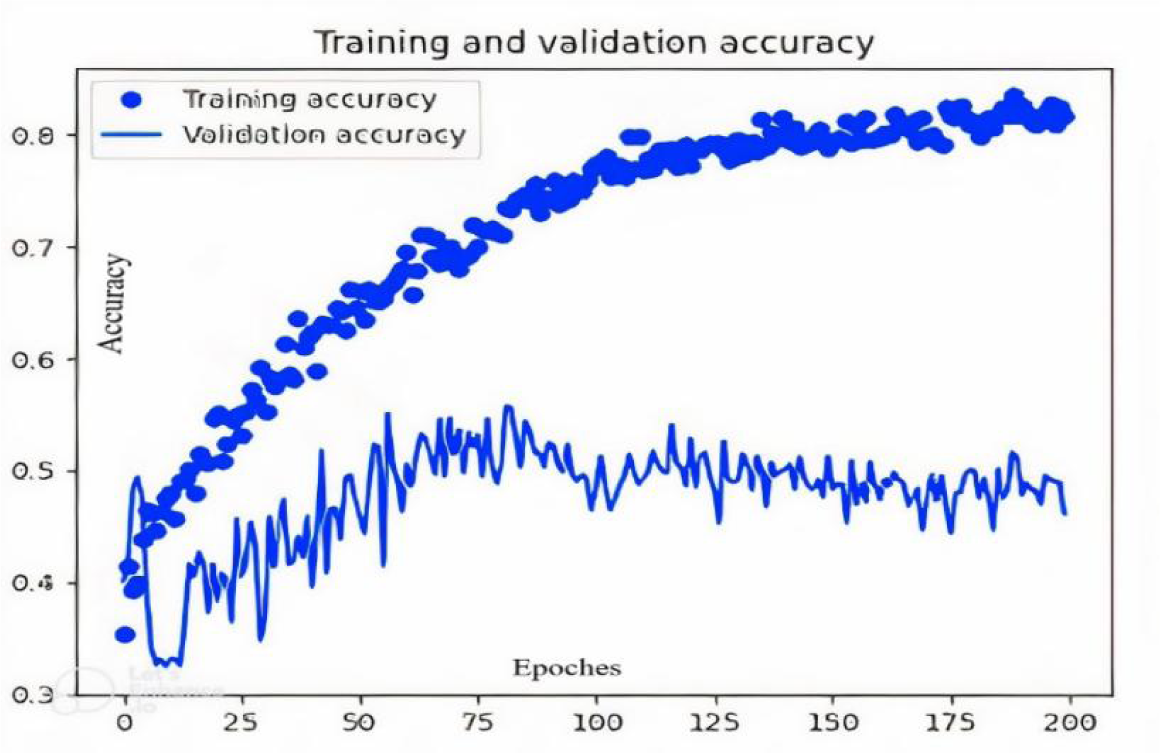
Learning rate 0.01 Accuracy 84.00%

**Figure 7.**
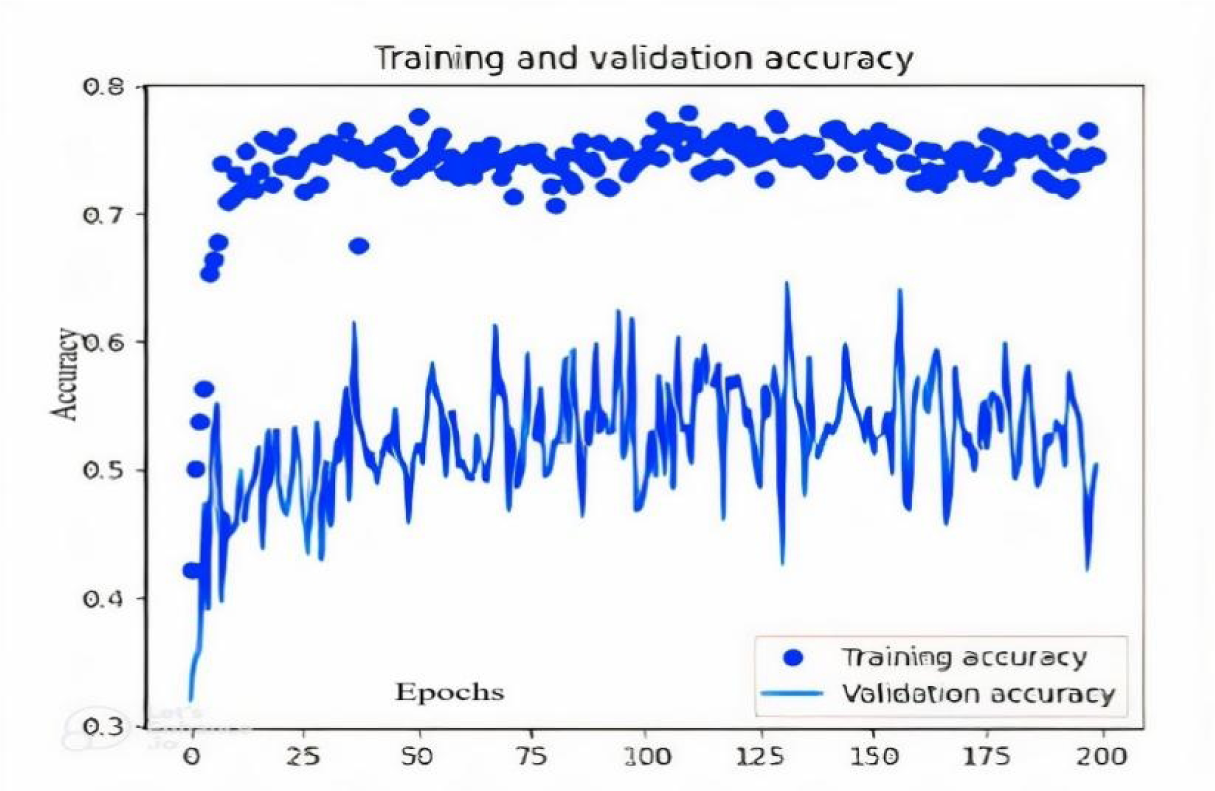
Learning rate 0.001 Accuracy 78.04%

**Figure 8.**
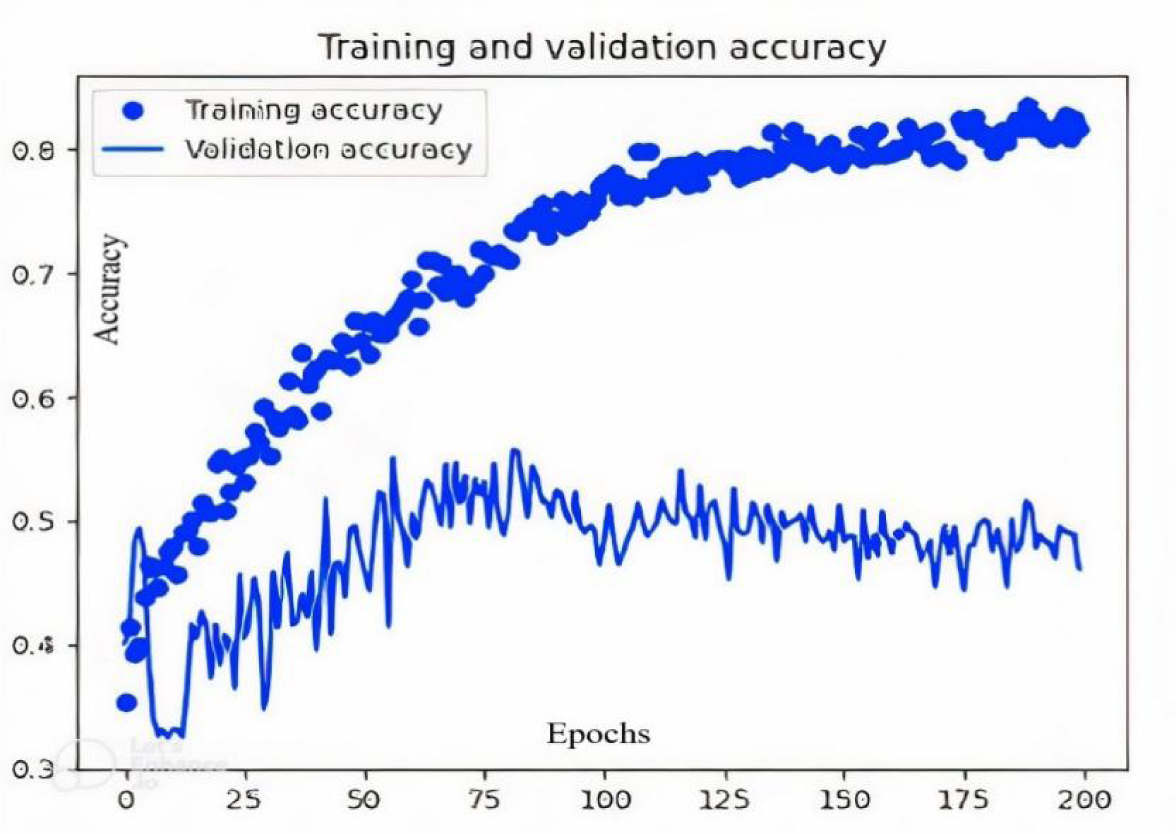
Optimizer Adam; Accuracy 84.64%

**Figure 9.**
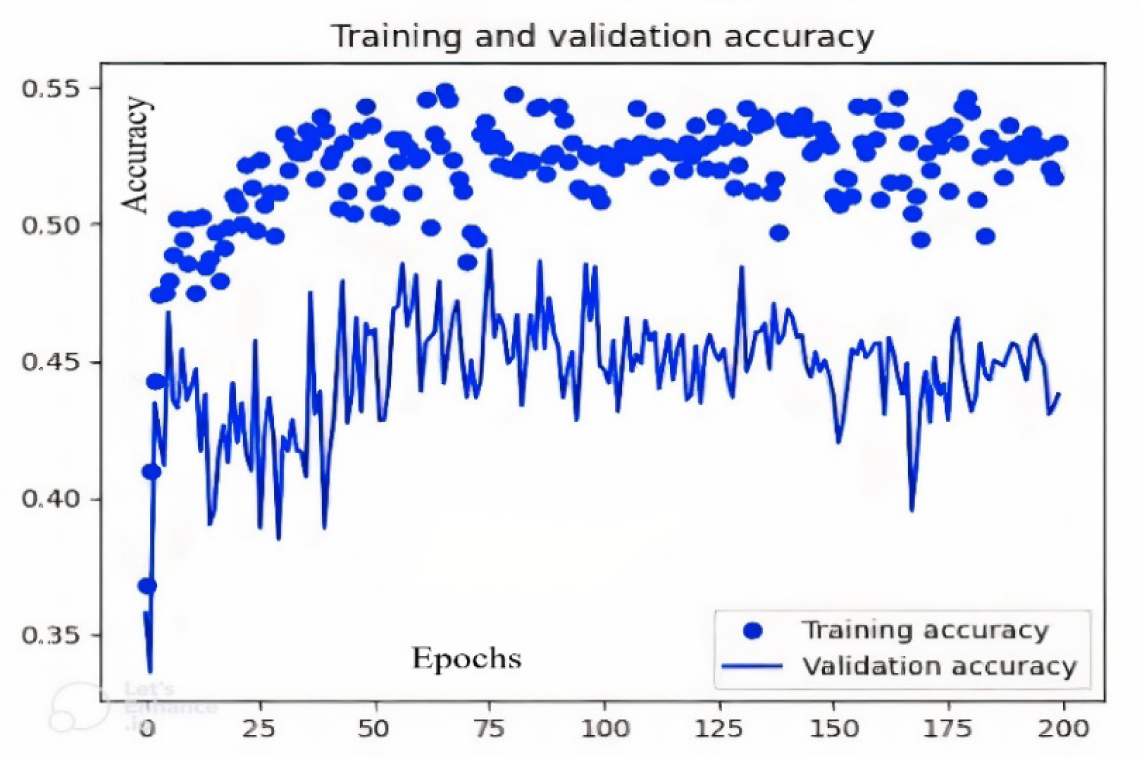
Optimizer Adadelta; Accuracy 54.92%

**Figure 10.**
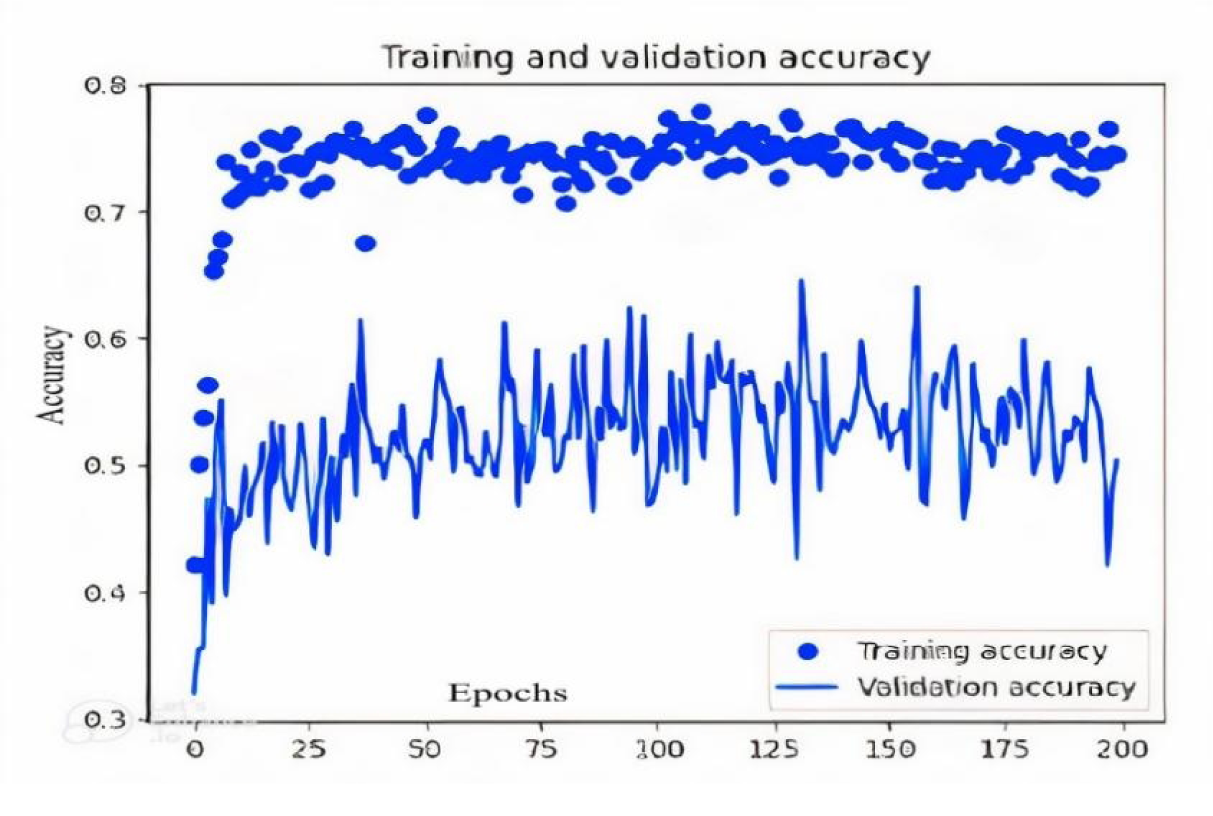
Batch-size 16 Accuracy 79.04%

**Figure 11.**
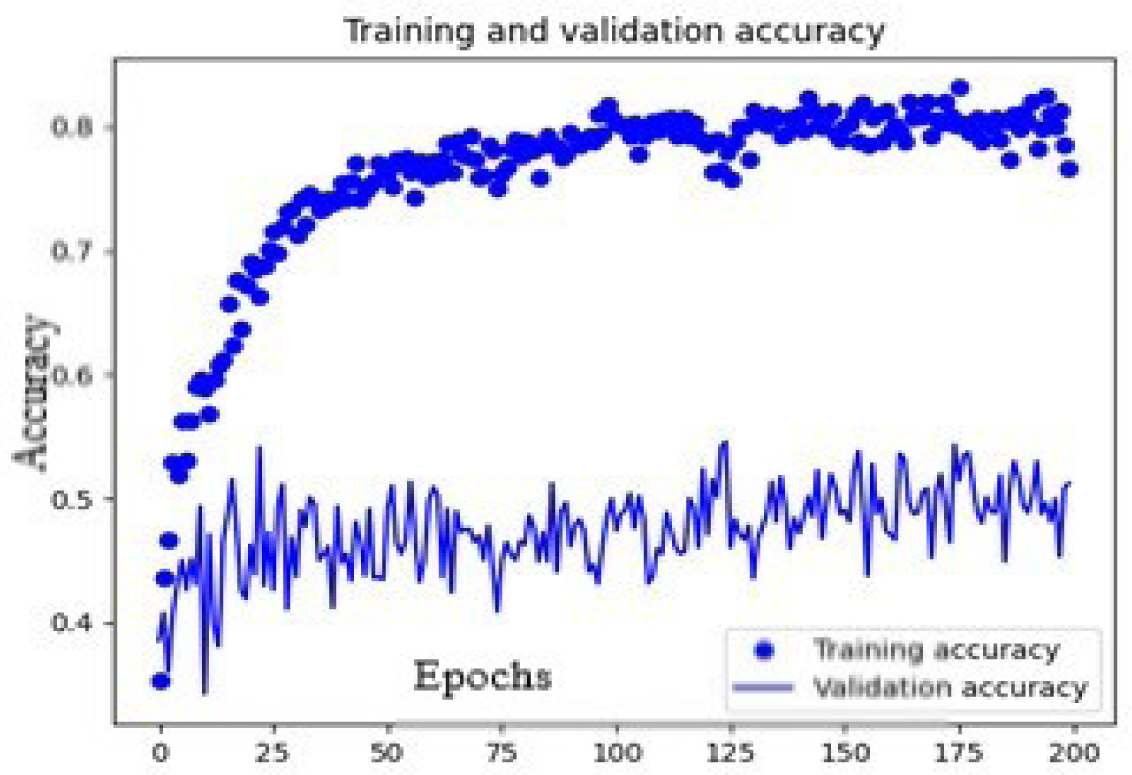
Batch-size 32 Accuracy 84.64%

We classified each dataset of repository into 3 classes i.e., Class 0 with no defects, Class 1 with one instance of defect, and Class 2 with instances of 2 or more than 2 numbers of defects.

### 3.2 Pre-Processing

Firstly, noise is eliminated from the dataset to normalize data. After that, we resolve the distribution gap between source and target dataset. Lastly, we solve the issue of class imbalance by using CTGAN Synthesizer. We get normalized data after all these steps to further use it for our experimentation. We are going to elaborate on each step of data preprocessing in detail. The values of features are different in updated repository (2021) in contrast to PROMISE repository (2020) on which previous researches [19] have been performed.

#### 3.2.1 Noise

We removed noise by eliminating the redundant data which is present in the PROMISE dataset so that defects are predicted accurately with higher accuracy by our trained model.

##### Method to Perform Experiment

We performed the following steps to remove noise from dataset:

- Input file in CSV format
- Eliminate insignificant columns
- Traverse each record of the dataset and select the duplicated rows and eliminate all duplicated records from the dataset.

Mu1tipe instances have been found which are duplicated in datasets. For example, Xerces-1.3 has duplicated rows which are considered as noise

#### 3.2.2 Distribution Gap

In the further step, we resolved the gap in distribution of data present in the dataset.

##### Method to Perform Experiment

Adversarial learning method is utilized to solve the data distribution gap problem. Firstly, we merged all the versions of datasets i.e., all versions of Xerces datasets as one file and we repeated the process for all datasets. We combined all available versions to get a higher volume of data for training purposes. We got 10 different datasets at the end. The following steps were performed:

- Take Source CSV File as input.
- Make a File which consist of column and traverse the loop to access each column.
- Min and max numbers are calculated, and their range values are listed for each and every column.
- Eliminate those data values which are outside of the min-max range.
- As a result, both the “source” and “target” dataset are in the same shape.
- Result prior and after covering the gap of data distribution is displayed in following figures

The distribution gap of the “dit” and “wmc” features of the camel-1.0 and xalan-2.4 dataset can be seen in the above graphs. Adversarial learning is used to cover this gap. We selected one dataset as the “source” and the other dataset as a “target”. Values that are outside of the min-max threshold value were assumed as outliers and we completely removed them to resolve the gap issue. The following is the figure after removing gap of distribution of data:

#### 3.2.3 Class Imbalance

Lastly, we balanced classes of the dataset through “CTGAN Synthesizer”.

##### Method to Perform Experiment

The following steps were performed to solve the class imbalance problem:

- Outliers are removed from the PROMISE repository, we then used CTGAN Synthesizer to generate synthetic clones for all source and target datasets from real data.
- During hyper-parameter tuning epoch was set to 100 since CTGAN Synthesizer generated the nearest clone data
- Select columns as parameters to train the CTGAN Synthesizer model.
- We generated synthetic data of all minority classes up to 966 instances since camel-1.6 has the highest number of instances among all the datasets.
- Datasets contain different numbers of instances in datasets which are shown in below table. Below tables display before and after experiment’s result.

CTGAN Synthesizer is used to make synthesis data to balance classes. Highest number of instances in any of the dataset are 966 so we produced all the dataset instances equal to 966. The class imbalance issue is resolved by generating the data on same pattern as dataset pattern which is learned by CTGAN Synthesizer

When we comprehensively analyzed the classes of dataset, it was apparent that many classes have less data which would degrade performance of prediction model. To solve the issue class zero and class one are taken as it is but merge the classes which have defects between 2 to 63 into Class 2. In this we get maximum data for training our model. As we have a maximum of 966 classes, 322 classes are generated for each of the bug class zero, class one, and class two. We can use data for the next step of experiment after balancing the classes.

### 3.3 Hybrid feature selection

We used HFS to select important features from the dataset. Hybrid methods are a good way to leverage the strengths of two algorithms instead of single method to select the most important features. “Filter & Wrapper Methods” and “Embedded and Wrapper Methods” are types of HFS available in literature [27]. “Embedded and Wrapper” method is chosen for experiment because Filter methods may not be able to search for the optimal subset of features sometimes, they consider each feature independently and does not check for feature dependencies on the other hand wrapper method is able to provide the best feature set [30]. Embedded methods select features according to their importance. [19] Selected optimal features set using a non-hybrid feature selection approach i.e., XGBoost. Whereas we used a HFS approach by combining wrapper method and embedded method. We chose the Random Forest algorithm as an embedded method. Random Forest is successful because it provides in general a good predictive performance and low over-fitting [31]. It is easy to find out the importance of the feature and then wrapper method is applied to rank the selected significant features. “Recursive Feature Elimination Cross-Validation” algorithm is used as a wrapper method in order to get our optimal feature subset. This method eliminates feature only one feature at each step instead of removing all features which is costly in terms of computational time. Basic RFECV works according to algorithm mentioned in referenced article [32]

There are 20 features in the dataset as discussed earlier. [19] Identified optimal feature set which is completely different in comparison to our significant feature selected though hybrid approach. Each\dataset has a different number of features selected through HFS according to their significance and relevance to particular dataset and these selected features plays most significant role in the results. Below Table 4 displays features selected for each dataset.

**Table 1:**
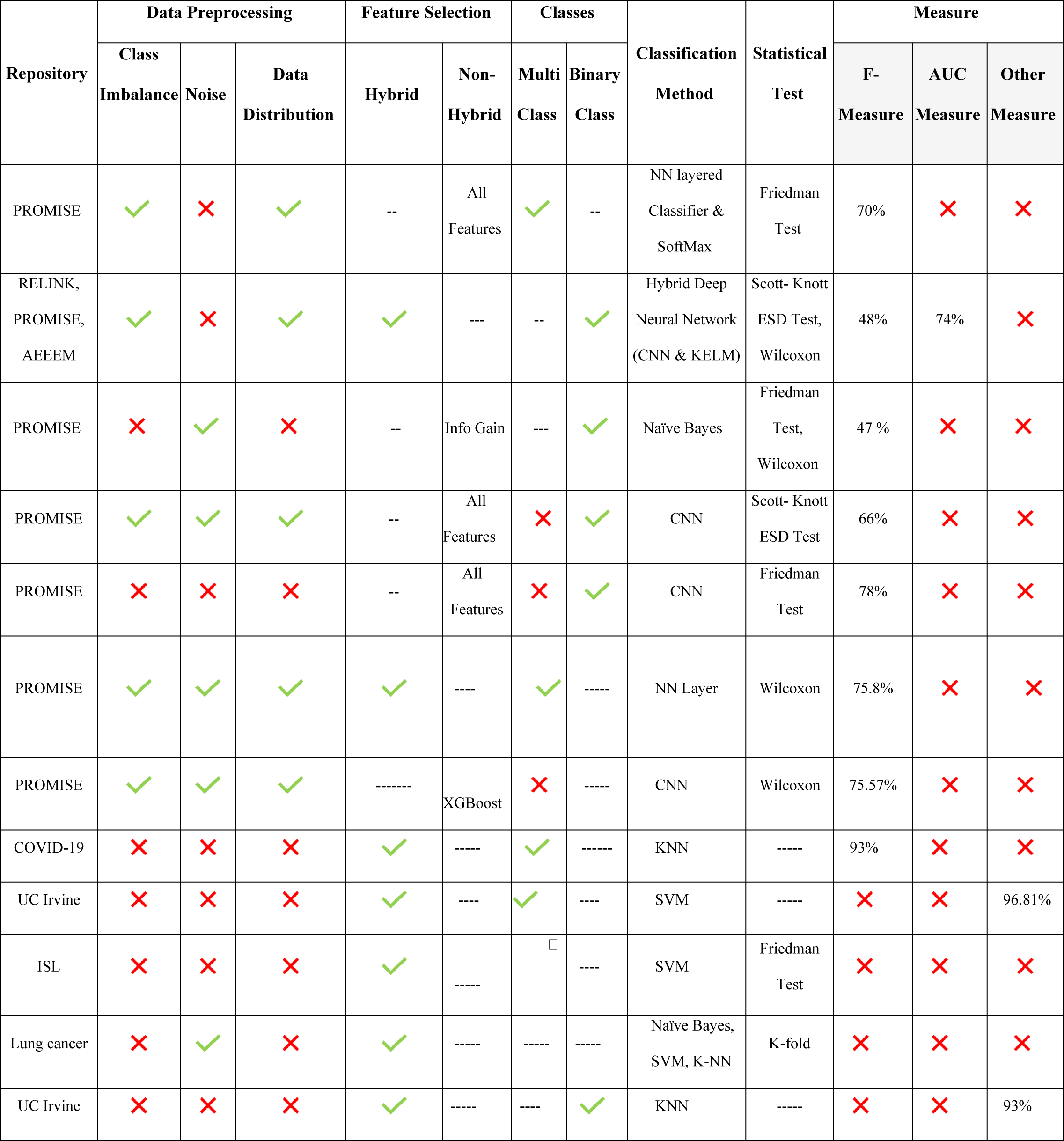
Summary table of literature review.

**Table 2.** Number of instances of datasets before resolving the imbalance of classes.

| Dataset | Number of Features | Total Instances |
| --- | --- | --- |
| "Ant 1.7" | 24 | 746 |
| "Camel 1.0" | 24 | 340 |
| "Camel 1.6" | 24 | 966 |
| "Jedit-3.4" | 24 | 273 |
| "Jedit-4.2" | 24 | 368 |
| "Log4j-1.1" | 24 | 110 |
| "Lucene-2.0" | 24 | 196 |
| "Poi-2.0" | 24 | 315 |
| "Synapse-1.0" | 24 | 158 |
| "Synapse-2.0" | 24 | 257 |
| "Velocity-1.6" | 24 | 230 |
| Xalan-2.4 | 24 | 724 |
| Xerces-1.2 | 24 | 441 |
| Xerces-1.3 | 24 | 454 |

**Table 3.** Number of instances in each dataset after balancing classes.

| Dataset | Number of Features | Total Instances |
| --- | --- | --- |
| “Ant 1.7” | 24 | 966 |
| “Camel 1.0” | 24 | 966 |
| “Camel 1.6” | 24 | 966 |
| “Jedit-3.4” | 24 | 966 |
| “Jedit-4.2” | 24 | 966 |
| “Log4j-1.1” | 24 | 966 |
| “Lucene-2.0” | 24 | 966 |
| “Poi-2.0” | 24 | 966 |
| “Synapse-1.0” | 24 | 966 |
| “Synapse-2.0” | 24 | 966 |
| “Velocity-1.6” | 24 | 966 |
| “Xalan-2.4” | 24 | 966 |
| “Xerces-1.2” | 24 | 966 |
| “Xerces-1.3” | 24 | 966 |

**Table 4.**
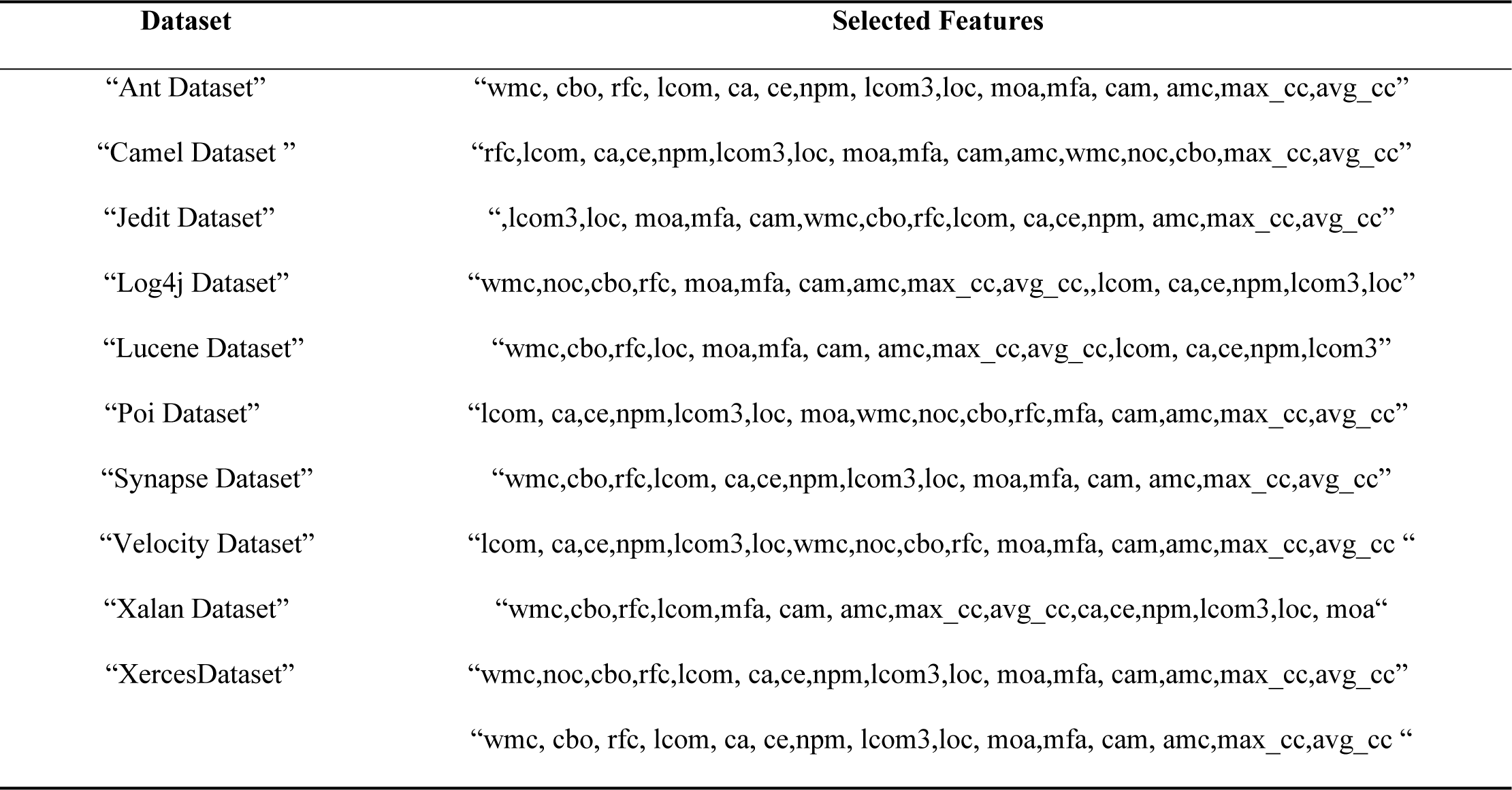
Significant features selection through HFS.

| Dataset | Selected Features |
| --- | --- |
| “Ant Dataset” | “wmc, cbo, rfc, lcom, ca, ce,npm, lcom3,loc, moa,mfa, cam, amc,max_cc,avg_cc” |
| “Camel Dataset ” | “rfc,lcom, ca,ce,npm,lcom3,loc, moa,mfa, cam,amc,wmc,noc,cbo,max_cc,avg_cc” |
| “Jedit Dataset” | “,lcom3,loc, moa,mfa, cam,wmc,cbo,rfc,lcom, ca,ce,npm, amc,max_cc,avg_cc” |
| “Log4j Dataset” | “wmc,noc,cbo,rfc, moa,mfa, cam,amc,max_cc,avg_cc,,lcom, ca,ce,npm,lcom3,loc” |
| “Lucene Dataset” | “wmc,cbo,rfc,loc, moa,mfa, cam, amc,max_cc,avg_cc,lcom, ca,ce,npm,lcom3” |
| “Poi Dataset” | “lcom, ca,ce,npm,lcom3,loc, moa,wmc,noc,cbo,rfc,mfa, cam,amc,max_cc,avg_cc” |
| “Synapse Dataset” | “wmc,cbo,rfc,lcom, ca,ce,npm,lcom3,loc, moa,mfa, cam, amc,max_cc,avg_cc” |
| “Velocity Dataset” | “lcom, ca,ce,npm,lcom3,loc,wmc,noc,cbo,rfc, moa,mfa, cam,amc,max_cc,avg_cc “ |
| “Xalan Dataset” | “wmc,cbo,rfc,lcom,mfa, cam, amc,max_cc,avg_cc,ca,ce,npm,lcom3,loc, moa“ |
| “XercesDataset” | “wmc,noc,cbo,rfc,lcom, ca,ce,npm,lcom3,loc, moa,mfa, cam,amc,max_cc,avg_cc” |
|  | “wmc, cbo, rfc, lcom, ca, ce,npm, lcom3,loc, moa,mfa, cam, amc,max_cc,avg_cc “ |

### 3.4 Dataset Division into Source and Target

We classified all source datasets through all relevant target datasets into multiple classes i.e., 0, 1 and 2. Since some of the datasets had minority instances, we therefore merged all the versions of same dataset into one CSV file. We tested “target” datasets against these merged “source” dataset to assess the accuracy of prediction model.

### 3.5 Classification

In this section we constructed the classification model which uses CNN model and SoftMax as final output layer. Following CNN structure has been used: Firstly, an input layer then three convolutional layers after that three non-linear activation functions are added then followed by a flatten layer, a fully connected layer and SoftMax layer as final layer. The classifier gets the instances as training data, passes through each layer and produce output which is a class probability. Our classifier classified output into three Class zero, Class one, and Class two. We evaluated the performance of model by taking each of the dataset as a “source” and “target” as well and repeated the entire process for each dataset.

### 3.6 Model Tuning

Our model is adjusted by changing epochs, learning rate, batch size, and optimizer to evaluate the model to a possible extent. It increased our understanding that which set of parameters setting gives the best performance of the proposed model.

### 3.7. Research Methodology

Our research type is explanatory and the discourse of our research is CPDP. Quantitative research methods are used because of numeric nature of datasets. We used datasets of PROMISE as input to train our model. We have selected significant features through HFS approach and evaluated its impact on prediction accuracy of defects for multi-class in cross datasets.

#### 3.7.4 Research Method

Experimental research method is used to investigate the cause and effect between dependent and independent variables for accuracy of prediction model. It is important to predict defect in their early phase by using existing non-defective datasets as reference. Defects are going to be predicted on early stage by testing our trained model on the target datasets. Our aim is selection of significant features through HFS approach and evaluate its impact on defects prediction accuracy for multi-class in cross projects, Source and target datasets of PROMISE repository are object of study in the experiment. Prediction Accuracy is the dependent variable of our research. Our research questions, hypothesis and experimental design of dependent and independent variable(s) in experimentation are unlike designed previously [19]. Wilcoxon test is used for defect prediction to validate results of research. We will be addressing the following research question:

**RQ1: Evaluate impact of significant feature selection through hybrid approach i.e., Random Forest & Recursive Feature Elimination Cross Validation upon defect prediction in Cross Project i.e., promise repository for multi class?**

- **Null Hypothesis (H0):** Significant feature selection through Hybrid technique i.e., Random Forest & Recursive Feature Elimination Cross Validation) has no impact on accuracy of CPDP i.e., promise repository for multi-class.
- **Alternate Hypothesis (H1):** Significant feature selection through Hybrid technique i.e., Random Forest & Recursive Feature Elimination Cross Validation) has impact on accuracy of CPDP i.e., promise repository for multi-class.

Our research has two manipulative independent variables i.e., All Features set and Selected Features Set through HFS i.e., Random Forest and Recursive Feature Elimination Cross Validation. Our research has one dependent variable i.e., Accuracy (AUC measure). One factor 2 treatment is used as our research design. Factor is going to be considered as design method and there are two Treatments: All features and selected features through HFS.

## 4 Result and Discussion

In this section, answer of research question and result analysis are discussed.

### 4.1 Hyper-Parameter Tuning

Optimizer, learning rate, and batch size hyper parameters are tweaked to see on which setting prediction model performs better.

#### 4.1.1 Learning Rate

We applied hyper-parameter tuning in terms of learning rate by considering the learning rate as 0.001 and 0.01. We considered Camel as the target dataset and Ant as source dataset. The best training and validation accuracy we achieved is by considering the learning rate as 0.01. The following figures show training and validation accuracy rates by considering epochs on the x-axis and accuracy on the y-axis.

#### 4.1.2 Optimizer

We applied hyper-parameter tuning by selection optimizers as Adam and Adadelta. We considered Camel as the target dataset and Ant as source dataset. The best training and validation accuracy we achieved is by considering an optimizer as adam. The following figures show training and validation accuracy rates by considering epochs on the x-axis and accuracy on the y-axis for different optimizers.

#### 4.1.3 Batch Size

We applied hyper-parameter tuning in terms of batch size by considering batch sizes as 16 and 32. We considered camel as the target dataset and ant as source dataset. The best training and validation accuracy we achieved is by considering batch size as 32. The following figures show training and validation accuracy rates by considering epochs on the x-axis and accuracy on the y-axis

From the above visualization, it is obvious that the optimal batch size is 32 for the dataset so, for all the datasets in the experiment we kept the “batch size” as 32.

### 4.2 Experimental Configuration

Experimental set up of hyper parameter values of CNN are unlike in previously [19] since we have experimented various combinations of optimizer i.e., Adam and Ada delta and the batch size on 16 and 32 and then in the end variations in learning rate i.e., 0.0001 and 0.01. We added two convolutional layers, leakyRelu layers, max pooling layers, dropout layers, and one flatten and one dense layer in architecture of CNN

### 4.3 Confusion Matrix

Confusion Matrix is a performance measurement used to define prediction model’s performance in terms of accuracy. Confusion matrix’s values demonstrate predictions of classes made by prediction model. In Figure 12, x-axis shows predicted values and y-axis shows true values. Out of 322 number of 0 classes, 241 are classified as zero class, 69 are classified as 1 class and 12 are classified as 2 class which means that 241 instances are classified correctly, and 81 instances are not correctly classified.

**Figure 12.**
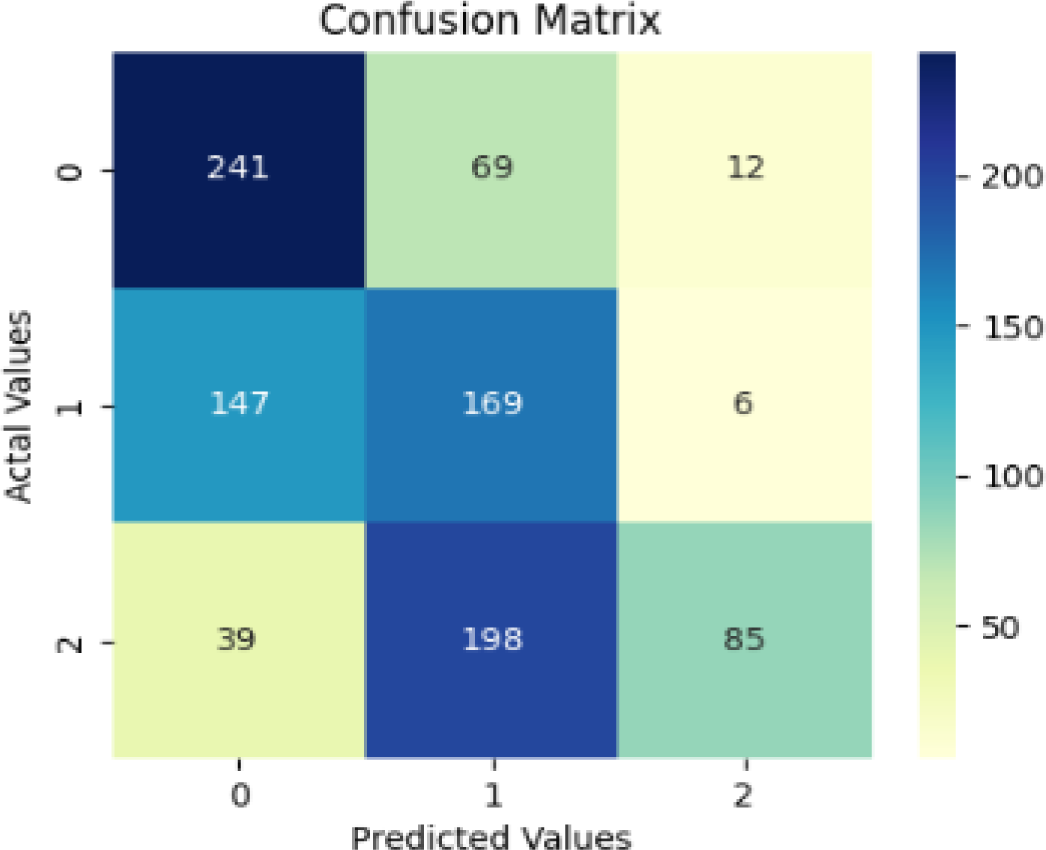
Confusion matrix while taking camel-1.0 (target) and ant (source) dataset

**RQ1: Evaluate the impact of significant feature selection through a hybrid approach i.e., Random Forest & Recursive Feature Elimination Cross Validation upon defect prediction in Cross Project i.e., promise repository for multi-class.**

We used projects of the PROMISE repository to perform an experiment and evaluated the results. Firstly, we pre-processed the data of source and target datasets. PROMISE repository consists of 10 datasets of which 4 datasets i.e., Camel, Jedit, Synapse, and Xerces have 2 versions each. We removed outlier values that are outside the range of minimum and maximum values of source and target datasets. After that, for better prediction accuracy, we merged versions of the same dataset to expand the total number of instances in that dataset so there are a high number of real data instances in comparison to synthetically generated clone data. The dataset is classified into 3 classes i.e., Class 0 with no defects, Class 1 with one instance of the defect and Class 2 with instances of 2 or more than 2 numbers of defects. This class imbalance issue is resolved by generating a synthetic data clone for each class using CTGAN Synthesizer. After that, the HFS approach is used to select significant features which played an important role in defect prediction. Then, we used CNN classifier and added SoftMax as final layer in model. AUC measure is used to evaluate the accuracy of proposed prediction model.

Firstly, results are displayed for every target and source dataset in AUC measure considering all features regardless of their insignificance.

The above Table 5. show the results of our experiment, we achieved an average of 72% AUC for all datasets when we used all feature for a model regardless of their insignificant in the above table dashes in the row show the similar source and target datasets which be within project defect prediction that’s why we are excluding them as we are experimenting for CP.

**Table 5:** Result of experiment in AUC % for each dataset considering all features.

| Target | Source Dataset |  |  |  |  |  |  |  |  |  |
| --- | --- | --- | --- | --- | --- | --- | --- | --- | --- | --- |
| Dataset | Ant<br>(Avg) | Camel<br>(Avg) | Jedit<br>(Avg) | Lucene<br>(Avg) | Poi<br>(Avg) | Log4j<br>(Avg) | Velocity<br>(Avg) | Synapse<br>(Avg) | Xalan<br>(Avg) | Xerces<br>(Avg) |
| Ant(Avg) | --- | 76 | 72 | 69 | 73 | 67 | 68 | 74 | 72 | 71 |
| Camel(Avg) | 76 | --- | 74 | 69 | 69 | 68 | 70 | 73 | 73 | 78 |
| Jedit(Avg) | 73 | 74 | --- | 69 | 66 | 67 | 65 | 71 | 72 | 70 |
| Lucene(Avg) | 68 | 70 | 67 | --- | 65 | 63 | 64 | 64 | 67 | 69 |
| Poi(Avg) | 70 | 75 | 66 | 62 | --- | 59 | 64 | 65 | 67 | 68 |
| Log4j(Avg) | 74 | 75 | 68 | 65 | 64 | --- | 64 | 68 | 71 | 76 |
| Velocity(Avg) | 69 | 76 | 66 | 60 | 64 | 60 | --- | 65 | 67 | 66 |
| Synapse(Avg) | 67 | 70 | 66 | 60 | 61 | 59 | 60 | --- | 62 | 67 |
| Xalan(Avg) | 70 | 73 | 70 | 64 | 68 | 62 | 64 | 69 | --- | 71 |
| Xerces(Avg) | 76 | 78 | 74 | 69 | 71 | 78 | 69 | 73 | 75 | 78 |

Now, we are going to display results for each and every target and source dataset of the repository and the results in AUC measure while considering significant features selected in our HFS approach.

The above Table 6 shows the results of our experiment in that we achieved higher defect prediction accuracy which is 78% average of all datasets in terms of AUC measure. There is a significant difference in prediction accuracy when considering all features of the dataset is an average of 72%. It clearly shows that features selected by our HFS are all significant features and played a significant in high defect prediction accuracy. The maximum accuracy we achieved in term of AUC is for **Camel** as a source dataset because this dataset and have maximum number of instances to train classifier for better prediction.

**Table 6:** Datasets which have high AUC % Accuracy considering selected features.

| Target | Source Dataset |  |  |  |  |  |  |  |  |  |
| --- | --- | --- | --- | --- | --- | --- | --- | --- | --- | --- |
| Dataset | Ant<br>(Avg) | Camel<br>(Avg) | Jedit<br>(Avg) | Lucene<br>(Avg) | Poi<br>(Avg) | Log4j<br>(Avg) | Velocity<br>(Avg) | Synapse<br>(Avg) | Xalan<br>(Avg) | Xerces<br>(Avg) |
| Ant(Avg) | --- | 84 | 81 | 76 | 78 | 82 | 78 | 75 | 78 | 84 |
| Camel(Avg) | 84 | --- | 84 | 78 | 84 | 85 | 84 | 78 | 80 | 86 |
| Jedit(Avg) | 78 | 82 | --- | 74 | 74 | 80 | 74 | 74 | 78 | 82 |
| **Lucene(Avg)** | --- | 77 | 77 | --- | 73 | --- | --- | --- | --- | 77 |
| **Poi(Avg)** | 76 | 79 | 76 | --- | --- | 75 | --- | --- | 76 | 79 |
| **Log4j(Avg)** | --- | 76 | 75 | --- | --- | --- | --- | --- | --- | 76 |
| **Velocity(Avg)** | --- | 78 | 76 | --- | --- | 75 | --- | 68 | --- | 78 |
| **Synapse(Avg)** | 77 | 81 | 79 | 74 | 73 | 77 | 73 | --- | 77 | 81 |
| **Xalan(Avg)** | 78 | 83 | 81 | 75 | 75 | 81 | 75 | --- | --- | 83 |
| **Xerces(Avg)** | 79 | 86 | 82 | 77 | 76 | 84 | 76 | 75 | 79 | --- |

In the above Table 7, the minimum accuracies we achieved in terms of AUC measure are by considering Log4j as a source dataset because we have a limited number of instances in each version. Due to this issue of limited number of instances we have to train the CTGAN to make a synthetic clone of data which results in better prediction accuracy for classifier.

**Table 7:** Datasets which have low AUC % Accuracy considering selected features.

| **Target** | **Source Dataset** |  |  |  |  |  |  |  |  |  |
| --- | --- | --- | --- | --- | --- | --- | --- | --- | --- | --- |
| **Dataset** | **Ant** | **Camel** | **Jedit** | **Lucene** | **Poi** | **Log4j** | **Velocity** | **Synapse** | **Xalan** | **Xerces** |
|  | (Avg) | (Avg) | (Avg) | (Avg) | (Avg) | (Avg) | (Avg) | (Avg) | (Avg) | (Avg) |
| **Ant(Avg)** | --- | --- | --- | --- | --- | --- | --- | --- | --- | --- |
| **Camel(Avg)** | --- | --- | --- | --- | --- | --- | --- | --- | --- | --- |
| **Jedit(Avg)** | --- | --- | --- | --- | --- | --- | --- | --- | --- | --- |
| **Lucene(Avg)** | 72 | --- | --- | --- | 70 | --- | 70 | 67 | 72 | -- |
| **Poi(Avg)** | --- | --- | --- | --- | --- | --- | 72 | 69 | --- | --- |
| **Log4j(Avg)** | 70 | --- | --- | 71 | 68 | --- | 68 | 67 | 70 | --- |
| **Velocity(Avg)** | 72 | --- | --- | 72 | 72 | -- | --- | 68 | 72 | --- |
| **Synapse(Avg)** | --- | --- | --- | -- | --- | --- | --- | --- | --- | --- |
| **Xalan(Avg)** | --- | --- | --- | --- | --- | --- | --- | 70 | --- | --- |
| **Xerces(Avg)** | --- | --- | --- | --- | --- | --- | --- | --- | --- | --- |

**Table 8:** Test statistics.

|  | P-Value | Accepted |
| --- | --- | --- |
| **Test Result** | 0.0032 | H1 |

As evident from our experimental results, significant feature are selected from hybrid approach which played important role in better defect prediction accuracy in term of AUC for multi-class in cross projects for promise repository which validate our H1 hypothesis i.e., “Significant feature selection through Hybrid technique i.e., Random Forest & Recursive Feature Elimination Cross Validation) has impact on defect prediction accuracy in cross project i.e., promise repository for multi-class “.

### 4.4 Research Validation

Wilcoxon statistical testing is used to validate results. Hypothesis for Wilcoxon test is either H0 when two sample groups are not significantly different from each other or H1 when two sample groups are significantly different from each other. In Wilcoxon test p-value is equivalent to 0.05. Acceptance condition is that “if the p-value is greater than 0.05, then H0 is accepted else H1 is accepted”.

For Wilcoxon test, we used accuracy values from experiment while considering all features of dataset as 1^st^ group and accuracy values which are obtained from experiment while considering selected feature through HFS approach as 2^nd^ group. As a result, we achieved a p-value of 0.0032 which is less than 0.05. It proves that sample distributions are not equal and approach of HFS through Random Forest and Recursive Feature Elimination Cross Validation does have a significant impact on the CPDP for multi-class problem.

## 5 Threat to Validity

Following Internal, external, and construct validity are three categories of possible threats to the validity which are pointed out in this research:

### 5.1 Optimal Hyper Parameter Setting

We tried different values of hyper parameters to optimize prediction accuracy of CNN model. However, it is possible to further explore variation in values and combination of hyper parameters to come up with optimal hyper parameter setting which may further improve prediction accuracy.

### 5.2 Internal Validity

How authentic are defects reported in PROMISE repository (2021). Since we worked on the undynamic features as written in the PROMISE repository.

### 5.3 Construct Validity

AUC measure is used to measure accuracy. It has been extensively used to measure accuracy of defect prediction models used in previous studies [5][8][9], so we believe that construct validity should be acceptable.

## 6 Conclusion

Cross-project defect prediction (CPDP) is a significant way of defect identification in the project. During experimentation upon 14 projects of PROMISE repository which is widely used in different field applications In Cross-project defect prediction, we extract knowledge from the source project and apply that learned knowledge to predict labels for the target dataset. However, the model performance is susceptible to significant feature selection. HFS can play a significant role in achieving high prediction accuracy. Exploratory Data Analysis proves that all versions of PROMISE repository are multi class. We selected significant features through HFS algorithms i.e., Random Forest, Recursive Feature Elimination Cross- Validation. We used CNN with SoftMax as the last layer to classify our results. After experimentation, we used Wilcoxon test to validate our results. As the result, our alternate hypothesis H1 “HFS technique (Random Forest & Recursive Feature Elimination Cross Validation) has an impact on CPDP accuracy for multi-class using PROMISE repository.” was accepted when p values evaluated as 0.0032 for Wilcoxon test.

## Author Contributions

Inception of Research, S.G. and R.B.F.; data curation, S.G. and R.B.F; formal analysis, M.A., S.G. and R.B.F.; funding acquisition, M.A., G.S. and A.Q.; Results and analysis, S.G. and R.B.F.; Proposed methodology, S.G.; Experimental Setup S.G.; Result validation, R.B.F., M. A., G. S. and A.A.; writing—original draft, S.G.; review and editing M.A., G.S., A.A., A. Q. and R.B.F. All authors have read and agreed to the published version of the manuscript.

## Conflicts of Interest

None.

## Funding

The research is funded by the Department of Computer Science, Zarqa University, Zarqa 13116 Jordan.

## Data Availability

Data repository used for experimentation is PROMISE 2021 which is publicly available at https://doi.org/10.6084/m9.figshare.13536506.v1

## Notes

### Competing Interest Statement

The authors have declared no competing interest.

## References

1. Hall, T., Beecham, S., Bowes, D., Gray, D., & Counsell, “A Systematic Literature Review on Fault Prediction Performance in Software Engineering” IEEE Transactions on Software Engineering, vol. 38, no. 6, pp. 1276–1304, 2012.

2. Nawaz, S., Zai, A., Imtiaz, S., & Ashraf, H, “Systematic literature review causes of rework in GSD,” Int. Arab J. Inf. Technol, vol. 19, no. 1, pp. 97–109, 2022.

3. Wang, S., Liu, T., Nam, J., & Tan, L, “Deep Semantic Feature Learning for Software Defect Prediction, “IEEE Transactions on Software Engineering, vol. 46, no. 12, pp. 1267–1293, 2020.

4. Menzies, T., Milton, Z., Turhan, B., Cukic, B., Jiang, Y., & Bener, A., “Defect prediction from static code features: current results, limitations, new approaches,” Automated Software Engineering, vol. 17, no. 4, pp. 375–4, 2010

5. Ma, W., Chen, L., Yang, Y., Zhou, Y., & Xu, B, “Empirical analysis of network measures for effort-aware fault-proneness prediction,” Information and Software Technology, vol. 69, pp. 50–70, 2016.

6. S. Herbold, A. Trautsch and J. Grabowski, “A Comparative Study to Benchmark Cross-Project Defect Prediction Approaches,” in IEEE Transactions on Software Engineering, vol. 44, no. 9, pp. 811–833, 1 Sept. 2018.

7. Kondo, M., Bezemer, C.-P., Kamei, Y., Hassan, A. E., & Mizuno, “The impact of feature reduction techniques on defect prediction models,” Empirical Software Engineering, vol. 24, no. 4, pp. 1925–1963, 2019.

8. Zhu, K., Ying, S., Zhang, N., & Zhu, D, “Software defect prediction based on enhanced metaheuristic feature selection optimization and a hybrid deep neural network,” Journal of Systems and Software, vol. 180, pp. 111026, 2021.

9. Li, J., He, P., Zhu, J., & Lyu, M. R, “Software Defect Prediction via Convolutional Neural Network,” 2017 IEEE International Conference on Software Quality, Reliability and Security vol. QRS, pp. 318–328, 2017.

10. Qiu, S., Xu, H., Deng, J., Jiang, S., & Lu, L, “Transfer Convolutional Neural Network for Cross-Project Defect Prediction,” Applied Sciences, vol. 9, no. 13, pp. 2660, 2019.

11. Wang, A., Zhang, Y., Wu, H., Jiang, K., & Wang, “Few-Shot Learning Based Balanced Distribution Adaptation for Heterogeneous Defect Prediction,” IEEE Access, vol. 8, pp. 32989–33001, 2020.

12. Ren, Y., Zhao, P., Sheng, Y., Yao, D., & Xu, “Robust SoftMax Regression for Multi-class Classification with Self-Paced Learning,” Proceedings of the Twenty-Sixth International Joint Conference on Artificial Intelligence, pp. 2641–2647, 2017.

13. Liu, Weiyang, Wen, Yandong, Yu, Zhiding, Yang, Meng, “Large-Margin SoftMax Loss for Convolutional Neural Networks,” Proceedings of the 33rd International Conference on Machine Learning, New York, NY, USA, 2016.

14. Hosseini, S., Turhan, B., & Mäntylä, “A benchmark study on the effectiveness of search-based data selection and feature selection for cross project defect prediction,” Information and Software Technology, vol. 95, pp. 296–312, 2018.

15. Sun, Y., Jing, X.-Y., Wu, F., Li, J., Xing, D., Chen, H., & Sun, Y, “Adversarial Learning for Cross-Project Semi-Supervised Defect Prediction,” IEEE Access, vol. 8, pp. 32674–32687, 2020.

16. Sheng, L., Lu, L., & Lin, “An Adversarial Discriminative Convolutional Neural Network for Cross-Project Defect Prediction,” IEEE Access, vol. 8, 55241–55253, 2020.

17. Pan, C., Lu, M., Xu, B., & Gao, “An Improved CNN Model for Within-Project Software Defect Prediction,” Applied Sciences, vol. 9, no. 10,pp. 2138, 2019.

18. Jalil, Abeer, Rizwan Bin Faiz, Sultan Alyahya, and Mohamed Maddeh.. “Impact of Optimal Feature Selection Using Hybrid Method for a Multiclass Problem in Cross Project Defect Prediction,” Applied Sciences vol. 12, no. 23: 12167, 2022.

19. Noreen, Sundas, Rizwan Bin Faiz, Sultan Alyahya, and Mohamed Maddeh. “Performance Evaluation of Convolutional Neural Network for Multi-Class in Cross Project Defect Prediction,” Applied Sciences vol. 12, no. 23: 12269, 2022.

20. Shaban, W. M., Rabie, A. H., Saleh, A. I., & Abo-Elsoud, M, “A new COVID-19 Patients Detection Strategy (CPDS) based on hybrid feature selection and enhanced KNN classifier,” Knowledge-Based Systems, vol. 205, pp.106270, 2020.

21. Ahmed, N., Rafiq, J. I., & Islam, M, “Enhanced Human Activity Recognition Based on Smartphone Sensor Data Using Hybrid Feature Selection Model,” Sensors, vol. 20, no. 1, pp.317, 2020.

22. Almugren, N., & Alshamlan, H, “A Survey on Hybrid Feature Selection Methods in Microarray Gene Expression Data for Cancer Classification,” IEEE Access, vol. 7, pp. 78533–78548, 2019.

23. Bansal, S. R., Wadhawan, S., & Goel, “mRMR-PSA Hybrid Feature Selection Technique with a Multiobjective Approach for Sign Language Recognition,” Arabian Journal for Science and Engineering, vol. 47, no.8, pp. 10365–10380, 2022.

24. Shukla, A. K., Singh, P., & Vardhan, “A New Hybrid Feature Subset Selection Framework Based on Binary Genetic Algorithm and Information Theory,” International Journal of Computational Intelligence and Applications, vol. 18, no. 3, 2019.

25. Al-Tashi, Q., Abdul Kadir, S. J., Rais, H. M., Mirjalili, S., & Alhussian, H, “Binary Optimization Using Hybrid Grey Wolf Optimization for Feature Selection,” IEEE Access, vol. 7, pp. 39496–39508, 2019.

26. Prinzie, A., & van den Poel, D, “Random Forests for multiclass classification: Random MultiNomial Logit,” Expert Systems with Applications, vol. 34, no. 3, pp.1721–1732, 2008.

27. Ren, Y., Zhao, P., Sheng, Y., Yao, D., & Xu, Z, “Robust Softmax Regression for Multi-class Classification with Self-Paced Learning,” Proceedings of the Twenty-Sixth International Joint Conference on Artificial Intelligence, pp.2641–2647, 2017.

28. Aggarwal, Deepti, “Software Defect Prediction Dataset,” figshare. Dataset, 2021.

29. A. Jović, K. Brkić and N. Bogunović, “A review of feature selection methods with applications,” 38th International Convention on Information and Communication Technology, Electronics and Microelectronics (MIPRO), pp. 1200-1205, 2015.

30. Saeys, Y., Abeel, T., Van de Peer, Y, “ Robust Feature Selection Using Ensemble Feature Selection Techniques. In: Daelemans, W., Goethals, B., Morik, K. (eds) Machine Learning and Knowledge Discovery in Databases,” ECML PKDD 2008. Lecture Notes in Computer Science(), vol 5212, 2008.

31. Chen, RC., Dewi, C., Hua “Selecting critical features for data classification based on machine learning methods,” J Big Data vol. 7, pp.52 2020.

32. Younes Charfaoui, “Hands-on with Feature Selection Techniques: Hybrid Methods, Part 5: Combining filter, wrapper, and embedded feature selection methods”.

